# Active site assembly by SMG5 as a mechanism for SMG6 endonuclease licensing in nonsense-mediated mRNA decay

**DOI:** 10.64898/2025.12.27.695470

**Authors:** Enes S. Arpa, Michael Taschner, Mara De Matos, Stefanie Jonas, David Gatfield

**Author notes:** shared first authorship (Enes S. Arpa, Michael Taschner). e-mail addresses.

## Abstract

Nonsense-mediated mRNA decay (NMD) is a conserved eukaryotic surveillance pathway that eliminates transcripts containing premature termination codons (PTCs). While the recognition phase of NMD is well characterised, the effector step – how targeted mRNAs are nucleolytically degraded – remains incompletely understood. In metazoans, NMD employs an endonucleolytic route mediated by SMG6, a PIN-domain nuclease, alongside SMG5 and SMG7, which act downstream of PTC recognition. SMG5 has recently been proposed to license SMG6 activity, yet the molecular basis of this licensing has remained elusive. Here, we combine AlphaFold structural predictions with biochemical assays to investigate interactions among human SMG5, SMG6, and SMG7. Structural models predict a high-confidence interface between SMG5 and SMG6 PIN domains that forms a composite active site: a conserved SMG5 aspartate (D893) complements the SMG6 acidic triad to reinstate the canonical tetrad required for PIN-domain catalysis. *In vitro*, SMG6 alone exhibits weak endonucleolytic activity, which is enhanced ∼10-fold by SMG5 PIN. Mutational analyses confirm that conserved residues from both proteins are essential for this composite configuration. Our findings reveal that the SMG5 PIN domain, previously considered catalytically inert, plays a critical role in activating SMG6 by completing its active site. This work provides mechanistic insight into the SMG5-dependent licensing step and uncovers a composite PIN nuclease architecture at the heart of the metazoan NMD effector phase.

## Introduction

Nonsense-mediated mRNA decay (NMD) was first recognized more than three decades ago as a conserved eukaryotic mRNA surveillance pathway that selectively degrades transcripts containing premature termination codons (PTCs) [1]. Early genetic screens in *Saccharomyces cerevisiae* [2] and *Caenorhabditis elegans* [3] identified the core Upf (up-frameshift) and Smg (suppressor of morphogenesis on genitalia) factors, defining a pathway that is conserved from yeast to humans. Subsequent work across species has established the “rules of NMD” – i.e., the molecular features that determine whether a terminating ribosome triggers or escapes NMD recognition and activation. By contrast, the effector phase – how targeted transcripts are ultimately degraded – took longer to resolve. NMD was long thought to rely primarily on accelerated general exonucleolytic mRNA turnover rather than specialized nucleases [4, 5]. It is now clear, however, that NMD recruits both exonucleolytic and endonucleolytic decay mechanisms, with striking mechanistic divergence between yeast and metazoans [6].

In yeast, NMD relies predominantly on the canonical decapping-dependent 5’→3’ exonucleolytic decay pathway [7]. Once marked for NMD, transcripts are decapped by Dcp1/Dcp2 and rapidly degraded by the 5’→3’ exonuclease Xrn1, with the 3’→5’ exosome contributing to a lesser extent [4, 8]. This paradigm established NMD as a pathway that co-opts general mRNA decay machinery rather than deploying pathway-specific nucleases.

By contrast, the situation in metazoans is more complex. Two key discoveries reshaped current models: first, that NMD substrates in *Drosophila melanogaster* undergo endonucleolytic cleavage near the PTC [9], and second, that *Drosophila* and human SMG6 proteins harbour an active PIN (<u>Pi</u>lT <u>N</u>-terminal) endonuclease domain domain [10–12]. In the prevailing model, SMG6 – representing the first dedicated nuclease intrinsic to the NMD machinery – directly cleaves target mRNAs, generating unprotected 3’ and 5’ fragments that are subsequently cleared by XRN1 and the exosome.

In addition to SMG6, metazoan NMD employs SMG5 and SMG7, factors not found in *S. cerevisiae*. All three proteins act downstream of PTC recognition i.e. after deposition of the central NMD effector helicase UPF1 on the target mRNA has been triggered. They directly associate with different sections of UPF1 [13], with SMG6 engaging the UPF1 N-terminus, the helicase domain [14] and a helicase regulating domain [15], while the SMG5-SMG7 module recognises C-terminal (phospho-)residues [16]. Historically, SMG5-SMG7 was viewed primarily as a heterodimeric adaptor that recruits general mRNA turnover enzymes – including the CCR4–NOT deadenylase complex and decapping factors [17–19] – thereby providing an exonucleolytic route analogous to the yeast mechanism. Although early work suggested that SMG6-mediated endo-cleavage and SMG5-SMG7-mediated exonucleolytic decay act in parallel and redundantly on overlapping sets of transcripts [20], more recent evidence supports a linear pathway, in which SMG5-SMG7 function upstream of SMG6 activation [21–23]. In particular, SMG5 has been proposed to act as a licencing factor for SMG6, such that SMG6 is recruited to UPF1 but requires a SMG5-SMG7-dependent authorisation step to execute endonucleolytic cleavage [21]. The nature of this authorisation signal remains unknown and is particularly intriguing given that, unlike SMG6, neither SMG5 nor SMG7 carry known catalytic activity and are presumed to act as scaffold proteins.

PIN domain proteins constitute a widespread family of single-stranded RNA nucleases in prokaryotes and eukaryotes. The canonical PIN core (∼120-130 amino acids) adopts a conserved fold – a parallel β-sheet flanked by α-helices – that forms an active site in which typically four acidic residues coordinate one or more divalent metal ions (often Mg^2+^ or Mn^2+^) to activate water for nucleophilic attack and RNA cleavage [24, 25]. Early structural studies of the human (*Hs*) SMG6 PIN domain [10] identified three key aspartate residues (D1251, D1353, D1392) – a reduced acidic cluster that, although less common within the structurally flexible PIN family, can suffice to support metal-dependent endonucleolytic activity (e.g. [26]). Indeed, *in vitro* assays demonstrated activity of a recombinant wild-type SMG6 PIN domain but not a D1353A mutant [10], and in a mouse model, mutation of two of the three Asp residues yielded effective NMD loss-of-function in vivo [27], validating the critical role of the SMG6 aspartate triade. Intriguingly, SMG5 also carries a C-terminal PIN domain with high similarity to that of SMG6, yet retains only a single conserved Asp (D860 in *Hs*SMG5); consistent with an impaired catalytic site, the recombinant SMG5 PIN domain lacks detectable cleavage activity [10].

Here, we investigate the predicted interactions among human SMG5, SMG6 and SMG7. AlphaFold3 [28] structure predictions recover the known SMG5-SMG7 interaction via their N-terminal domains and, in addition, predict with high confidence a physical interface between the SMG5 and SMG6 PIN domains that would create a composite active site. In this arrangement, a conserved aspartate from SMG5 (D893) complements the SMG6 aspartate triade to yield the acidic quartet typical of canonical PIN nucleases. We validate these predictions using *in vitro* nuclease assays, showing that the SMG6 PIN domain alone has weak nuclease activity that is ∼10-fold enhanced in the presence of the SMG5 PIN domain. Mutational analyses further identify conserved residues required for activity of the composite PIN configuration. Together, our data provide mechanistic insight into the function of the previously presumed inactive SMG5 PIN domain and suggest a molecular basis for the hypothesised SMG5-dependent authorisation step – namely, complementation of the SMG6 active site.

## Results

### Structural models predict a conserved SMG5-SMG6 PIN domain interaction

To explore how the metazoan-specific NMD factors SMG5, SMG6, and SMG7 assemble, we performed AlphaFold3 predictions on the full-length human proteins (Figure 1 and SI Figures S1 and S2). The resulting high-confidence model faithfully recapitulates the known SMG5-SMG7 complex formed via their N-terminal 14-3-3-like/α-helical domains [18], as well as the established intramolecular contact between the 14-3-3-like and α-helical domain of SMG6 [13] (SI Figures S1A-B and S2A). Unexpectedly, however, the model simultaneously predicts a previously unrecognized complex between the C-terminal PIN domains of SMG5 and SMG6 (Figure 1B). The central interface is formed by interdigitating helices from both domains (SMG5 α4/5 and SMG6 α8; Figure 1D), with additional predominantly hydrophobic contacts contributed by N-terminal α-helical extensions (SMG5 α1 and SMG6 α1; Figure 1E, F). Moreover, a loop in SMG6 that links its 14-3-3-like and α-helical domains is predicted to fold back onto the central SMG6 PIN β-sheet, contributing an extra β-strand (SMG6 β1) that forms several backbone hydrogen bonds with the SMG5 PIN domain (Figure 1F). While the minimal PIN-PIN interface alone covers ∼70 Å^2^, these auxialiary contacts expand the putative interaction surface to ∼190 Å^2^. Importantly, this domain arrangement leaves the SMG6 PIN active site fully accessible for RNA binding and even suggests that SMG5 may contribute catalytically relevant residues to the ribonuclease centre (Figure 1C). To assess evolutionary conservation, we generated analogous predictions for *Drosophila melanogaster* (*Dm*, for SMG5 and SMG6, as no SMG7 is encoded in this organism [29]) and *Caenorhabditis elegans* (*Ce*, whose orthologous proteins are considerably shorter [18]). Both species yielded closely similar architectures, notably preserving the SMG5-SMG6 PIN-PIN interface and the auxiliary SMG5-6 α1 and SMG6 β1 contacts (SI Figures S1C-F, S2B-C and S3A-B). Consistently, sequence alignments and surface mapping revealed strong conservation of interface residues in both SMG5 and SMG6, as well as for the predicted SMG6 β1 strand (Figures 2B, C and SI Figure S3C).

**Figure 1.**
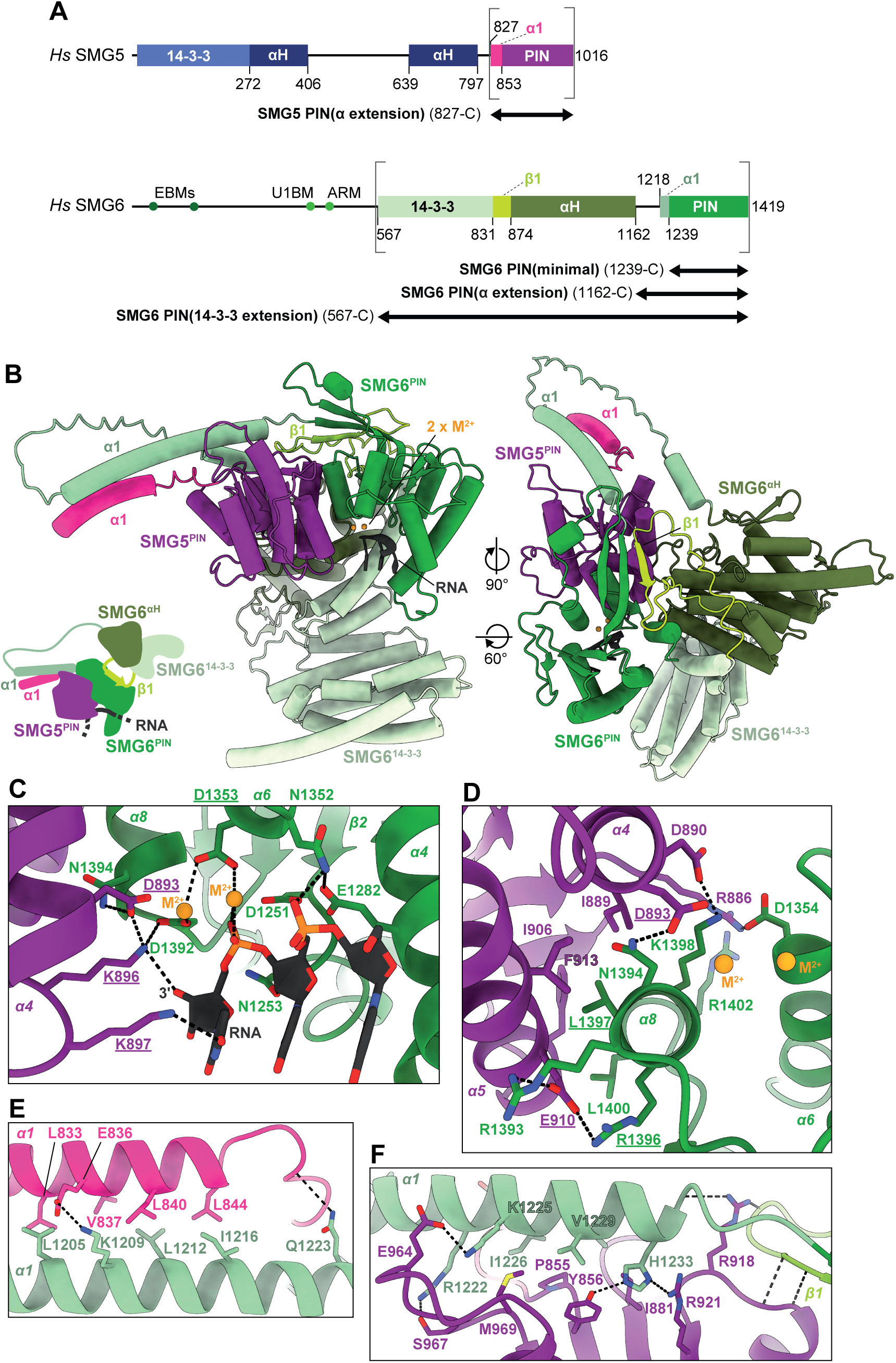
Structure models of SMG5 and SMG6 predict formation of a composite PIN domain. (A) Schematic domain architecture of *Hs*SMG5 and SMG6. SMG6 binding motifs for exon junction complex (EBMs 39-59 and 133-153) and UPF1 (U1BM 406-419 and ARM 445-465) are indicated by spheres. Constructs used in the study, i.e. SMG5 PIN(α extension) (827-C), SMG6 PIN(minimal) (1239-C), SMG6 PIN(α extension) (1162-C) and SMG6 PIN(14-3-3 extension) (567-C), are indicated at the bottom of each cartoon. The α-helices located N-terminally to the PIN domains that are predicted to interact are highlighted in colour and labelled α1. The β-strand, which is inserted in a loop between SMG6 14-3-3-like (14-3-3) and α-helical (αH) domain and which is predicted to intramolecularly complement the PIN-domain β-sheet is indicated as β1. (B) Model of the complex between *Hs*SMG5 PIN domain (purple, pink) and the folded domains of *Hs*SMG6 (shades of green) based on an AlphaFold3 model of the *Hs*SMG5-6-7 complex with U_6_-RNA and 2 Mg^2+^. Domains are coloured as in (A). RNA (grey) and metal ions (yellow spheres) in the active site are shown. Elements that contribute to the interaction i.e. α1-helices and β1-strand are highlighted and labelled. (C) Zoom into the proposed composite active site. Active site residues of SMG6 (green) and SMG5 (purple) are shown as sticks and labelled. Residues mutated in this study are underlined. Predicted positions of two bivalent cations (M^2+^) that are coordinated by the acidic residues, as well as three nucleotides of RNA substrate (dark grey) are also shown. Dotted lines indicate putative hydrogen bonds and salt bridges. (D) Zoom into the *Hs*SMG5PIN-SMG6PIN interface viewed from the bottom of the active site shown in (C). Residues of SMG5 (purple) and SMG6 (green) that are predicted to contribute to binding are shown as sticks and labelled. Residues mutated in the study are underlined. Dotted lines indicate putative hydrogen bonds or salt bridges. (E) Predicted interface between α-helical extensions (α1) of *Hs*SMG5 (pink) and SMG6 (light green) PIN domains. Interface residues are shown as sticks and labelled. (F) Predicted interface between SMG6 α1 (light green) and β1 with SMG5PIN (purple). Dotted lines indicate putative hydrogen bonds or salt bridges.

**Figure 2.**
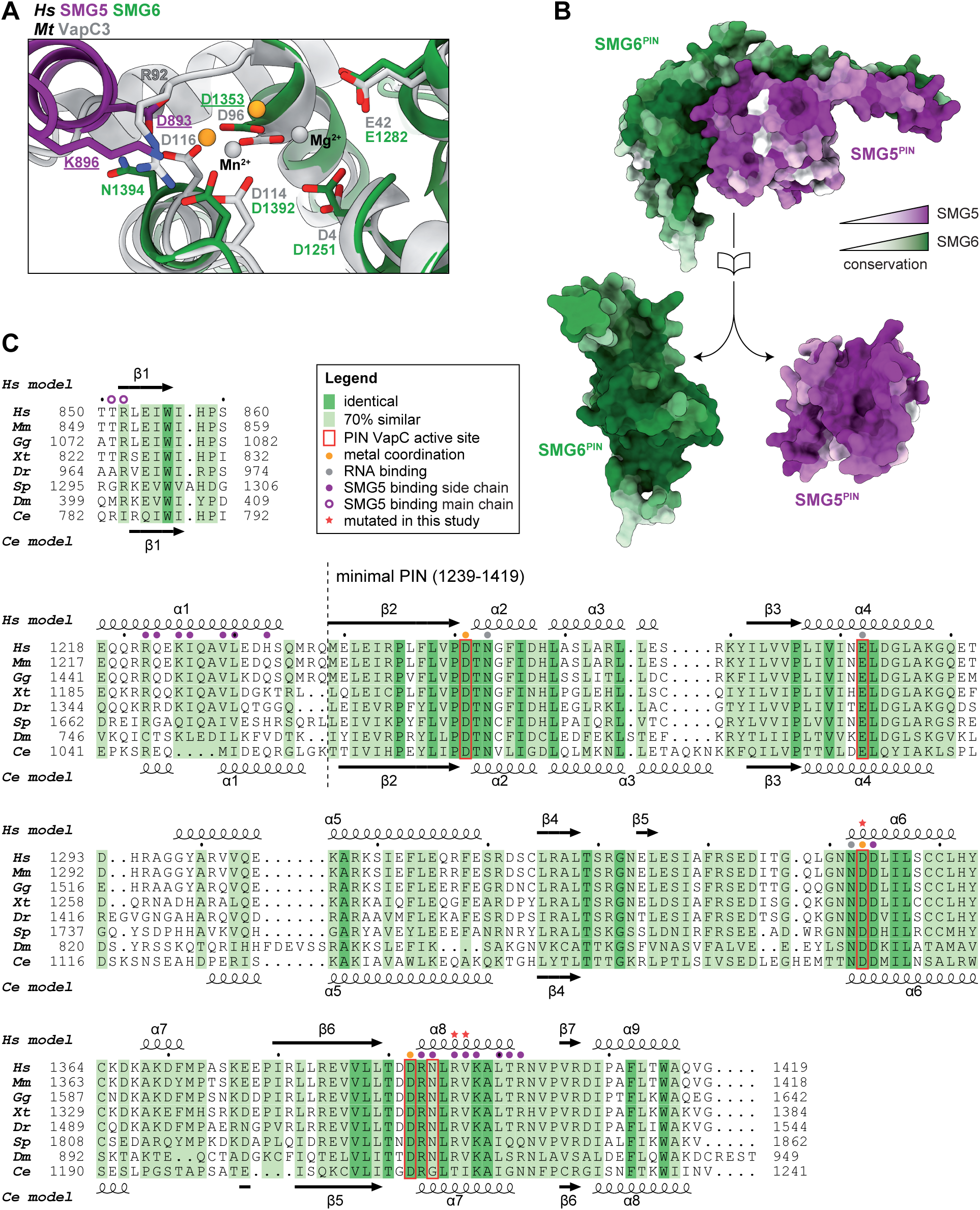
The composite SMG5-6 PIN active site and domain interface are evolutionarily conserved. (A) Superposition of the crystal structure of prototypical PIN domain family member *Mt*VapC3 (light grey, PDB ID 4CHG)[30] shows that it displays a similar arrangement of active site residues as the composite SMG5-6 PIN domain. Active site residues are shown as sticks and labelled. The two metal ions crystallized with VapC (Mn^2+^, Mg^2+^) are shown as grey spheres. (B) Surface conservation of the SMG5 (purple) and SMG6 (green) PIN domain interface, with increasing color darkness corresponding to increased conservation. (C) Sequence alignment of the extended SMG6 PIN domain including β1 strand and α1 helix. Orthologs from *Homo sapiens* (*Hs*), *Mus musculus* (*Mm*), *Gallus gallus* (*Gg*), *Xenopus tropicalis* (*Xt*), *Danio rerio* (*Dr*), *Strongylocentrotus purpuratus* (*Sp*, Purple urchin), *Drosophila melanogaster* (*Dm*) and *Caenorhabditis elegans* (*Ce*) are compared, and residues with complete conservation and 70% similarity are highlighted in dark and light green, respectively. Predicted secondary structure elements of *Hs* and *Ce* proteins are denoted above and below the alignment, respectively. Residues involved in the predicted SMG5 interface (purple), active site metal coordination (yellow) and RNA substrate binding (grey) are indicated by circles above the alignment. Red stars mark amino acids that were mutated in this study, while red borders mark residues that correspond to VapC active site components.

### Evolutionary conservation of residues forming a putative composite SMG5-SMG6 active site

The prediction that the SMG5 PIN domain sits in close proximity of the SMG6 PIN active site – potentially extending several side chains into its catalytic cavity – raises the possibility that SMG5 contributes directly to RNA cleavage. To assess the plausibility of this model, we compared the predicted SMG5-SMG6 active site configuration with canonical PIN domain family members. Previous bioinformatic analyses [24, 25] identified the SMG6 PIN domain as most closely related to the PIN domains of bacterial VapC toxins, which are highly active ribonucleases [26]. Structural superposition of the SMG5-SMG6 PIN complex with the crystal structure of *Mycobacterium tuberculosis* VapC3 [30]) (Figure 2A) revealed that the isolated SMG6 PIN lacks one of the four conserved aspartate residues that form the catalytic tetrad (*Mt*VapC3: D4, D96, D114, D116; *Hs*SMG6: D1251, D1353, D1392, **N**1394). These four Asp residues normally coordinate two divalent metal ions (Mg^2+^ or Mn^2+^ in many PIN domains) that activate a nucleophilic water molecule and stabilize the pentavalent transition state during phosphodiester hydrolysis. Notably, the absent fourth Asp is replaced by Asn in *Hs*SMG6 and *Dm*Smg6, and even by glycine in *Ce*SMG-6 (SI Figure S3A, B). Strikingly, association with SMG5 is predicted to supply a fully conserved aspartate (*Hs*SMG5 D893, *Dm*Smg5 D1050, *Ce*SMG-5 D464) positioned almost identically to the missing *Mt*VapC3 D116, thereby potentially complementing the SMG6 catalytic triad. In addition, two conserved Lys sidechains of SMG5 (*Hs*SMG5 K896, K897) are predicted to reach into the active site entrance in a position compatible with RNA binding; their position overlaps with an analogous Arg side chain in VapC3. Together the structural models, sequence, and evolutionary comparisons support the intriguing possibility that SMG5 and SMG6 cooperate to form a composite PIN-domain active site that is catalytically competent for RNA cleavage.

### Mn^2+^-dependent *in vitro* nuclease activity of SMG6 is enhanced by the SMG5 PIN domain

Based on the above structural predictions, we next sought to validate them experimentally. We first assessed the *in vitro* nuclease activity of the SMG6 PIN domain in the absence and presence of the SMG5 PIN domain. Previous work by Glavan *et al.* used a gel-based assay with a radioactively end-labelled U_30_ RNA oligonucleotide to measure SMG6 activity [10]. As a starting point, we developed an analogous fluorescence-based assay using a modified U_25_ RNA oligonucleotide protected against exonucleolytic degradation by deoxyU residues at both ends and carrying a 6-carboxyfluorescein (6-FAM) label at the 5′ end (Figure 3A). In the standard assay, the probe was incubated at a 1:1 molar ratio with recombinant, N-terminally tagged proteins in the presence of Mn^2+^, and substrate turnover was monitored after 45 min by 10% denaturing urea polyacrylamide gel electrophoresis (PAGE).

**Figure 3.**
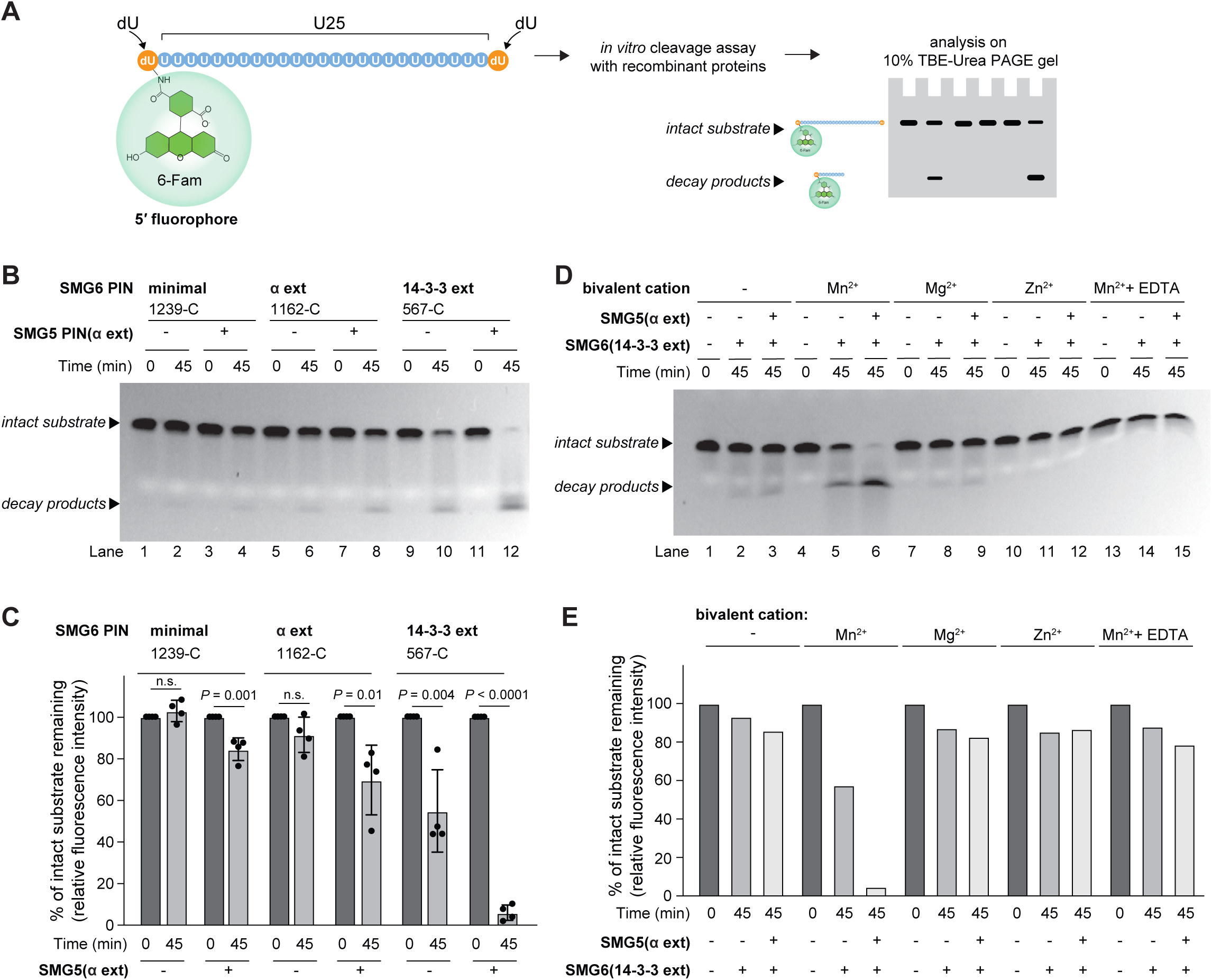
SMG5-PIN and SMG6-PIN together degrade RNA in a Mn^2+^ dependent manner. (A) Schematic of the gel-based *in vitro* RNA degradation assay using 5’-6FAM-conjugated RNA oligo (5’-6FAM-deoxyU-U_25_-deoxyU-3’). (B) Cleavage assay using 3 µM 5’-6FAM RNA oligo incubated with 3 µM of the indicated protein constructs in 5 µL of reaction buffer (100 mM NaCl, 5 mM MnCl_2_, 1 mM DTT, 10% glycerol, 25 mM Tris-HCl pH 7.5) for 45 min (or 0 min for controls) at 37 °C. Reactions were run on 10% PAGE-Urea gels. The bands corresponding to intact substrate and decay products are indicated. (C) Quantification of RNA gel degradation assay from (C) and from three additional replicates of identical experiments. The amount of remaining probe (upper band) with regard to the 0 min timepoint (=100%) was quantified. Data are plotted as means ± SD. *P*-values were calculated using two-tailed unpaired *t*-test. (D) Similar to experiment shown in panel (C), 3 µM of substrate and 3 µM of indicated protein con-structs were incubated with either 5 mM MnCl_2_, MgCl_2_, or ZnCl_2_ in a 5 µL reaction volume (buffer conditions: 100 mM NaCl, 1 mM EGTA, 1 mM DTT, 10% glycerol, 25 mM Tris-HCl pH 7.5). After 0 min and 45 min at 37 °C, reactions were analysed by PAGE. 30 mM EDTA (last 3 lanes) was used to show the specificity of Mn^2+^-mediated RNA degradation. (E) Quantification of degradation assay shown in (E).

To probe the predicted SMG5-SMG6 interaction, we produced three SMG6 PIN constructs of increasing length (Figure 1A and SI Figure S4): (i) a minimal PIN domain (aa 1239–1419); (ii) a short extended version (aa 1162–1419), termed SMG6 PIN(α ext), which includes the predicted α-helix (α1); and (iii) a long extended version (aa 567–1419), termed SMG6 PIN(14-3-3 ext), also comprising the 14-3-3-like and α-helical domains that, according to the structural models, contact the PIN domain through a short central β-strand (β1). The SMG5 construct used throughout (aa 827–1016, termed SMG5 PIN(α ext)) was slightly longer than that used by Glavan *et al.* (aa 853–1016; [10]), as it included the predicted α-helical extension (α1), which in the structural models forms stabilizing contacts with the SMG6 α1 helix to reinforce the PIN-PIN interface.

Side-by-side comparisons revealed that the SMG6 PIN(14-3-3 ext) construct combined with SMG5 PIN(α ext) exhibited the highest activity, achieving near-complete substrate degradation (∼95% turnover after 45 min; Figure 3B, lanes 11-12; Figure 3C). Notably, this SMG6 construct also displayed substantial basal activity in the absence of SMG5 (∼50% turnover; lanes 9-10). By contrast, combining SMG5 with shorter SMG6 constructs yielded progressively weaker activity: the minimal SMG6 PIN and SMG6 PIN(α ext) constructs turned over only ∼20% and ∼30% of substrate, respectively (lanes 1-8), and both constructs were essentially inactive without SMG5 (Figure 3C).

All assays were conducted in the presence of Mn^2+^, the preferred cofactor for many PIN domain nucleases. Using the SMG5 PIN(α ext)-SMG6 PIN(14-3-3 ext) combination, we verified metal ion specificity: substituting Mn^2+^ with Mg^2+^ or Zn^2+^ abolished cleavage, and chelation with EDTA blocked Mn^2+^-dependent degradation (Figure 3D, E).

In summary, SMG5 PIN markedly enhances SMG6 PIN nuclease activity *in vitro*, and structural extensions on SMG6 make substantial contributions to achieving maximal activity.

### Extended SMG5-SMG6 contacts can be functionally mimicked by covalent tethering

Our initial interpretation of the high activity observed with the structurally extended SMG6 construct was that the additional sequences stabilise a PIN-PIN interaction between SMG5 and SMG6 that would otherwise be weak. However, we could not exclude the possibility that these SMG6 extensions contribute directly to catalysis or RNA binding rather than acting solely as structural stabilisers.

To discriminate between these possibilities, we used the SMG5 PIN(α ext) and the SMG6 PIN(α ext) constructs, which exhibited low activity in our assays (Figure 3B, lanes 7-8; Figure 3C), and irreversibly linked them into a single polypeptide using the SpyTag/SpyCatcher system [31]. Remarkably, the covalently tethered SMG5-SMG6 pair displayed markedly higher activity than the corresponding non-tethered proteins. In fact, the SpyTag/SpyCatcher-linked SMG5 PIN(α ext)-SMG6 PIN(α ext) construct approached the activity of the unlinked SMG5 PIN(α ext)-SMG6 PIN(14-3-3 ext) proteins (Figure 4A, B). These findings indicate that the additional SMG6 sequences primarily act to stabilise SMG5-SMG6 association and become dispensable when physical proximity is enforced by other means. In our assay, this was achieved via covalent linkage; *in vivo*, an analogous effect may be mediated by co-recruitment of SMG5 and SMG6 to the common binding platform, phospho-UPF1.

**Figure 4.**
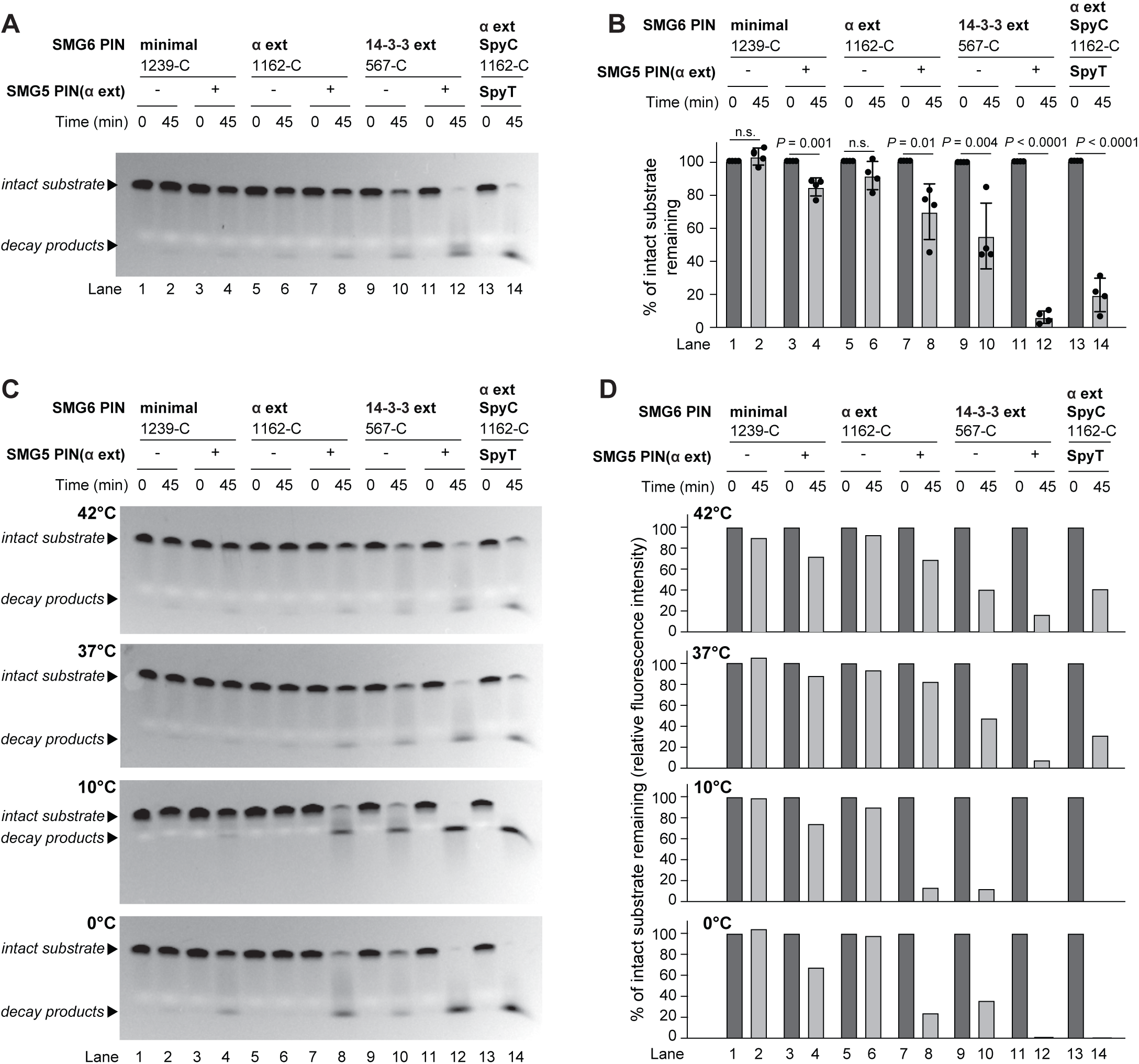
Covalent tethering and low temperature stabilize weak SMG5-SMG6 PIN domain interaction to enhance catalytic activity. (A) Cleavage assay using 3 µM 5’-6FAM RNA oligo incubated with 3 µM of the indicated protein constructs in 5 µL of reaction buffer (100 mM NaCl, 5 mM MnCl_2_, 1 mM DTT, 10% glycerol, 25 mM Tris-HCl pH 7.5) for 45 min (or 0 min for controls) at 37 °C. Reactions were run on 10% PAGE-Urea gels. Lanes 13 and 14 show the covalently tethered SMG5 PIN(α extension) and SMG6 PIN(α ex-tension), using the SpyTag/SpyCatcher strategy. (B) Quantification of RNA gel degradation assay from (A) and from three additional replicates of identical experiments. The amount of remaining probe (upper band) with regard to the 0 min timepoint (=100%) was quantified. Data are plotted as means ± SD. *P*-values were calculated using two-tailed unpaired *t*-test. Note that apart from the values in lanes 13 and 14 – showing the SpyTag/SpyCatcher-tethered SMG5-SMG6 pair – this graph reports on the same experimental data as in Figure 3 (B) for lanes 1-12. (C) As in (A), yet with experiments carried out at reaction temperatures of 42, 37, 10, or 0 °C. (D) Quantification of the assay shown in (C).

As a second, orthogonal test of the predicted weak interaction, we assessed the temperature-dependence of SMG5-SMG6 activity. Our rationale was that temperature exerts opposing effects on two key factors: enzymatic kinetics and protein-protein interaction. Lower temperatures generally slow catalytic turnover but can stabilise weak interfaces, whereas higher temperatures accelerate intrinsic reaction rates yet destabilise fragile interactions. Thus, if SMG5 and SMG6 interact weakly, we would expect increased activity at low temperature (due to improved complex stability) and reduced activity at high temperature, despite faster kinetics. Conversely, a strongly interacting dimer would be expected to show the opposite trend – slower activity at low temperature and faster at high temperature.

We tested this by performing the *in vitro* cleavage assay at four temperatures: 0 °C (on ice), 10 °C, 37 °C (as in Figure 3B), and 42 °C (proxy for upper physiological limit in humans). The results were unambiguous: low temperatures strongly enhanced substrate turnover by SMG6 variants that were nearly inactive at 37 °C. For example, the SMG6 PIN(α ext) together with SMG5 PIN(α ext) degraded only ∼30-40% of substrate at 37 °C–42 °C but achieved 80-90% turnover at 0 °C–10 °C (Figure 4C, lanes 7-8; Figure 4D). Additional observations from the temperature experiment included: (i) even the minimal SMG6 PIN acquired measurable activity with SMG5 at 0°C (30-40% turnover; lanes 3-4); and (ii) basal activity of SMG6 constructs alone also increased at low temperature – for instance, SMG6 PIN(14-3-3 ext) turned over ∼60% of substrate at 37 °C but 70-90% at 0 °C–10 °C (lanes 9-10). These temperature-dependent improvements may reflect enhanced RNA binding or improved structural stability and/or Mn^2+^ coordination at low temperature.

Together, these results demonstrate that the SMG5-SMG6 PIN domains form a weak, temperature-sensitive interaction that can be functionally stabilised either by extended structural elements or by enforced physical proximity. This supports a model in which auxiliary contacts on SMG6 reinforce the composite active site formed with SMG5.

### Quantitative comparison of nuclease kinetics through a real-time fluorescent cleavage assay

The gel-based endpoint assays above provided important insights into SMG6 nuclease activity and its modulation by SMG5, but they were unsuitable for kinetic analysis. To enable quantitative rate measurements, capture early reaction phases, and increase throughput, we developed a fluorescence-based real-time cleavage assay. This platform is particularly advantageous for analysing subtle effects of protein mutations on nuclease activity. We used a modified U_25_ RNA oligonucleotide substrate carrying a 6-FAM fluorophore at the 5′ end and a Black Hole Quencher 1 (BHQ1) at the 3′ end. In the intact substrate, BHQ1 quenches the 6-FAM signal, while cleavage separates the two moieties and results in increased fluorescence (Figure 5A).

**Figure 5.**
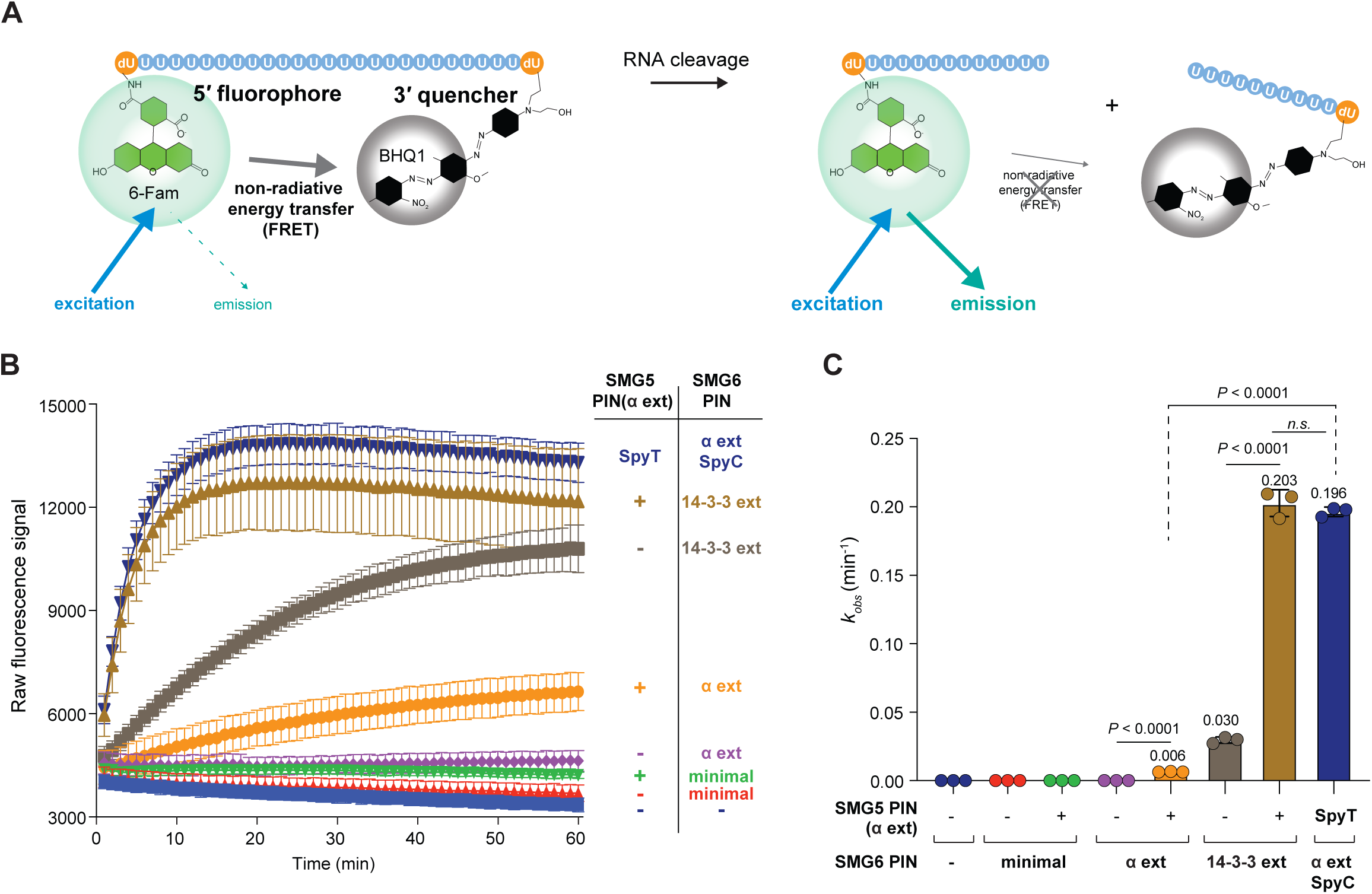
Real-time fluorescent cleavage assay reveals kinetic differences among SMG5-SMG6 constructs. (A) Schematic of the real-time fluorescent cleavage assay using the 5’-6FAM-dU-U_25_-dU-3’-BHQ1 substrate (termed 5’-6FAM-3’-BHQ1 in remainder of figure legends). (B) 3 µM 5’-6FAM-3’-BHQ1 oligo were incubated with 3 µM of the indicated protein constructs (or RNase-free water, as control) in 10 µL volume (reaction buffer: 100 mM NaCl, 5 mM MnCl_2_, 1 mM EGTA, 1 mM DTT, 10% glycerol, 25 mM Tris-HCl pH 7.5) for 60 min at 25 °C, with measurements at 1 min interval in a plate reader. Raw fluorescence values were plotted for each reaction (each con-dition run in triplicates) as medians ± SD. (C) Observed rate constants (*k_obs_*) calculated from each sample shown in (B). Data are plotted as means ± SD. The *P*-values were calculated using two-tailed unpaired *t*-test.

We benchmarked our assay using conditions similar to the gel-based assay. Reactions were initiated by protein addition, and fluorescence was recorded every minute for 60 min using a plate reader. The assay reproduced all preceding findings while enabling precise kinetic comparisons. SMG5 PIN(α ext) with SMG6 PIN(14-3-3 ext) and the SpyTag/SpyCatcher-tethered SMG5-SMG6 pair were highly active, reaching 50% substrate turnover in <5 min (Figure 5B). SMG6 PIN(14-3-3 ext) alone displayed clear activity but required >20 min to reach 50% turnover, whereas SMG5 PIN(α ext) with SMG6 PIN(α ext) showed only weak activity and all other constructs were inactive.

We fitted the fluorescence curves with a single-exponential model to derive apparent rate constants (*k_obs_*). The most active constructs yielded *k_obs_* values of ∼0.2 min^-1^, whereas SMG6 PIN(14-3-3 ext) alone was almost 7-fold slower (*k_obs_* ≈ 0.03 min^-1^) (Figure 5C).

In summary, this real-time assay provides fine-resolution kinetic measurements and a robust platform for dissecting SMG5/SMG6 function and mutant phenotypes.

### Mutational analysis validates SMG5-contributed active site residues and the SMG5-SMG6 interaction surface

To validate the structural model, we generated SMG5 and SMG6 mutants targeting two features: the composite active site (Figures 1C and 6A) and the predicted PIN-PIN interface (Figures 1D and 6B). First, we probed the composite active site. Recombinant SMG6 PIN(14-3-3 ext) carrying a D1353A mutation was nucleolytically inactive in all conditions, confirming the essential role of this catalytic residue (Figure 6A, C). On the SMG5 side, converting D893 to alanine abolished the ability of SMG5 PIN(α ext) to enhance SMG6-mediated cleavage, consistent with its proposed role in complementing the active site. Furthermore, mutating the conserved K896 and K897 – located near the composite active site and potentially involved in RNA binding – also eliminated SMG5-mediated stimulation. Analogous results were found for these mutants in the SpyTag/SpyCatcher setup (SI Figure S5A, B).

**Figure 6.**
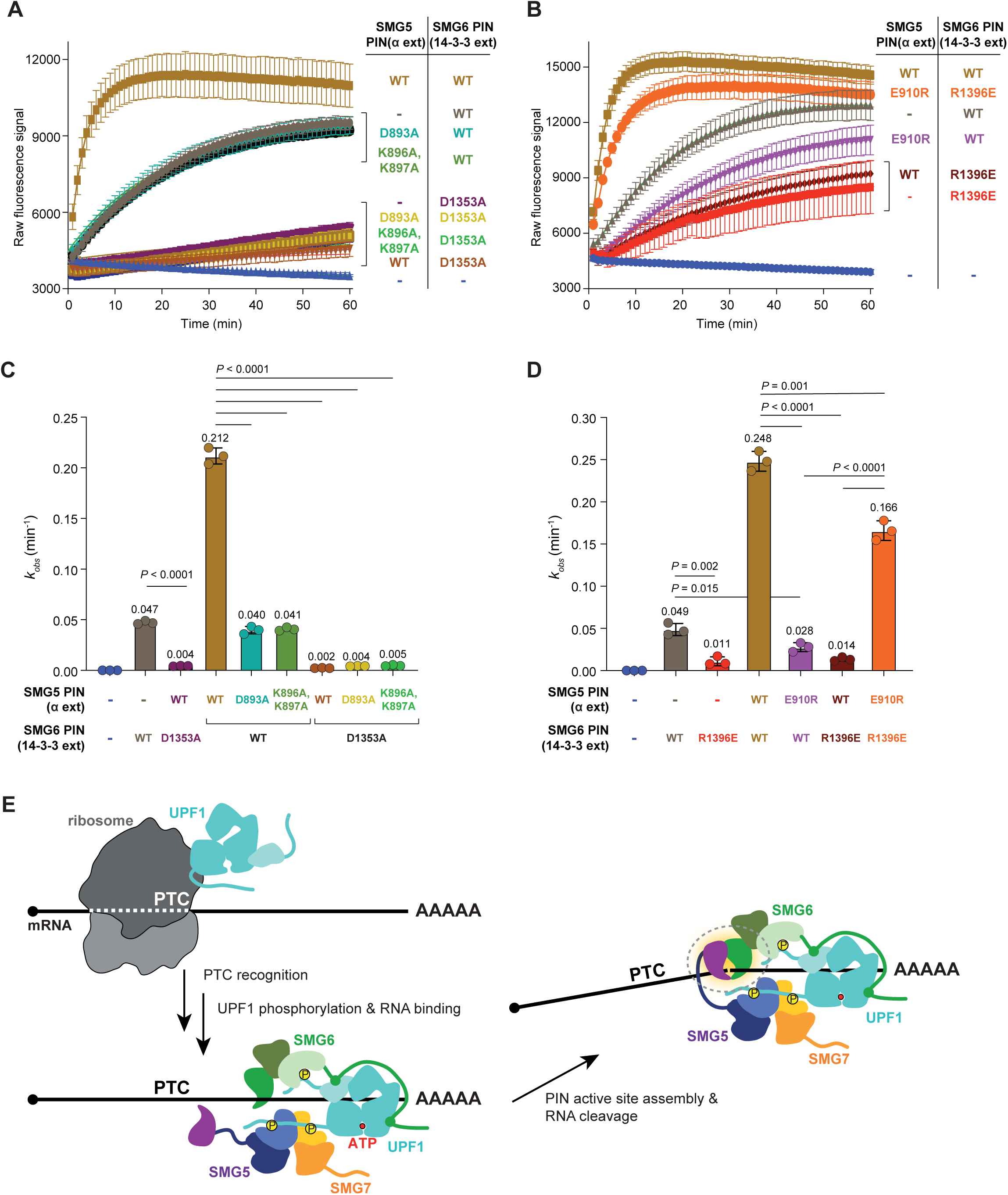
Mutational analysis confirms composite active site and PIN-PIN interface in SMG5-SMG6 complex. (A) 3 µM 5’-6FAM-3’-BHQ1 oligo were incubated with 3 µM of the indicated protein constructs that probe the catalytic site residues and its environment (SMG5 D893A, K896A, K897A and SMG6 D1353A) in 10 µL volume (reaction buffer: 100 mM NaCl, 5 mM MnCl_2_, 1 mM EGTA, 1 mM DTT, 10% glycerol, 25 mM Tris-HCl pH 7.5) for 60 min at 25 °C, with measurements at 1 min interval in a plate reader. Raw fluorescence values were plotted for each reaction (each condition run in tripli-cates) as averages with error bars corresponding to ± SD. (B) Identical to (A), but for mutants that probe the interaction interface (SMG5 E910R, SMG6 R1396E). (C) Observed rate constants (*k_obs_*) calculated from each sample shown in (A). Data are plotted as means ± SD. The *P*-values were calculated using two-tailed unpaired *t*-test. (D) As in (C), for data shown in (B). (E) Model of SMG5-dependent activation of SMG6 PIN nuclease through co-recruitment to phosphorylated UPF1 that is bound to PTC containing mRNA.

Next, we interrogated the predicted interaction surface. Structural models suggested an electrostatic contact between SMG5 E910 and SMG6 R1396 (Figure 1D). Disrupting this interaction by charge reversal (SMG5 E910R or SMG6 R1396E, together with a WT partner protein) greatly reduced activity (*k_obs_* ≈ 0.03 min^-1^ and 0.01 min^-1^, respectively), compared with that of the two WT proteins (*k_obs_* ≈ 0.25 min^-1^; Figure 6B, D). Strikingly, combining both mutants (SMG5 E901R + SMG6 R1396E) restored activity almost fully (*k_obs_* ≈ 0.17 min^-1^), directly validating the predicted electrostatic interaction. A second interface mutant, SMG6 V1397E, also prevented SMG5-mediated stimulation (SI Figure S5C-F). Interestingly, both SMG6 interface mutants reduced basal activity even in the absence of SMG5, conceivably because introducing a negative charge (glutamate) near the active site could perturb RNA binding.

In summary, these mutational analyses confirm the structural model of active site complementation between SMG5 and SMG6 and identify conserved residues critical for catalysis and PIN–PIN interaction.

## Discussion

The precise roles of SMG5, SMG6, and SMG7 in the effector phase of metazoan NMD have remained unclear for decades. While SMG6 was established as the pathway’s dedicated endonuclease, SMG5 and SMG7 were largely considered scaffolds that recruit general decay factors such as deadenylases and decapping enzymes. Our findings revise this view by demonstrating that SMG5 directly contributes to catalysis: its PIN domain complements residues missing from the SMG6 active site, generating a composite nuclease architecture. This mechanism explains why the isolated SMG6 PIN domain is unable to trigger mRNA decay *in vivo* even upon tethering [32], why SMG5 acts as a licensing factor for SMG6 [22, 23], and it provides a molecular basis for the proposed authorisation step in NMD [21]. Together with prior literature, our data supports a model in which the SMG5-SMG7 heterodimer is recruited efficiently to the phosphorylated C-terminus of UPF1 via its two adjacent 14-3-3-like domains [13, 16, 18, 33, 34], while SMG6 recognizes both the RNA-bound conformation of UPF1 and its phosphorylated N-terminus [13–15]. Only the avidity afforded by this combination of several low affinity interactions brings the two PIN domains of SMG6 and SMG5 into proximity to elicit stable active site formation and productive RNA cleavage (Figure 6E). The intramolecular connection between the SMG6 14-3-3-like domain and its PIN domain may help ensure that cleavage occurs *in cis*, in close proximity to the UPF1 RNA binding site and the PTC [12, 35].

Composite active sites are not unprecedented in the PIN domain superfamily. Structural studies of bacterial VapC toxins and other PIN-containing effectors have shown that dimerization configures catalytic grooves, often bringing together acidic residues from two domains [26, 36]. Similar principles underlie many PIN-domain and related nucleases, where catalysis is conditional upon oligomerization or conformational change. For instance, CRISPR-associated PIN enzymes can be activated by cyclic tetraadenylate (cA_4_)-dependent oligomerization that reconfigures active-site residues across different subunits [37]. Our work extends this paradigm to eukaryotic RNA surveillance, revealing that SMG5 and SMG6 cooperate to assemble a canonical PIN active site that neither domain can fully provide alone.

This conditional assembly offers an elegant regulatory strategy. Nuclease activation is contingent upon protein-protein interactions and co-recruitment to a common platform – phosphorylated [13, 16] and RNA-bound [15] UPF1 – ensuring tight spatial and temporal control during NMD. The requirement for SMG5-mediated complementation effectively establishes a checkpoint that licences SMG6 activity only within correctly assembled surveillance complexes. Such conditional activation also prevents ectopic SMG6 activity in the cytoplasm, safeguarding transcripts from unintended cleavage.

Evolutionary comparisons reinforce this model. *Drosophila melanogaster* lacks SMG7 but retains SMG5 and SMG6 [29], suggesting that the SMG5-SMG6 partnership represents the catalytic core of metazoan NMD. SMG7 may represent an additional, yet dispensable adaptor to enhance exonucleolytic decay or pathway regulation, whereas SMG5’s contribution to SMG6 activation appears indispensable. This observation underscores that SMG5 is not merely a scaffold but a critical licensing factor for SMG6 endonucleolytic activity.

Beyond clarifying SMG5’s role, these findings highlight composite PIN architectures as a recurrent evolutionary solution for licensing nuclease activity. Conditional assembly allows organisms to couple catalytic potency to molecular context, a principle that may apply broadly to RNA decay pathways and other nucleases. Future work should examine whether SMG7 contributes additional layers of regulation, how UPF1 phosphorylation dynamics influence composite active site formation, and whether similar composite architectures exist in other metazoan RNA surveillance systems. Understanding these mechanisms will not only refine models of NMD regulation but may also inform therapeutic strategies targeting aberrant RNA decay in genetic disease and cancer.

## Methods

### Cloning

Human *SMG5* and *SMG6* sequences were amplified by PCR from plasmids pcDNA3.1-MS2-HA-HsSmg5_T (Addgene #147552) [19] and pCIneo-λN-HA-HsSmg6_I (Addgene #146551) [38], or from HEK293 cell cDNA, and then cloned by Golden-gate cloning into low-copy plasmid pLIBT7 in fusion with N-terminal His-tag (for SMG5), N-terminal HisGST-tag (for SMG6), N-terminal SpyTag [31] and C-terminal His-tag (for SMG5) or N-terminal SpyCatcher [31] and C-terminal His-tag (for SMG6). For bacterial co-expression, SMG5 plasmids carried ampicillin resistance and SMG6 plasmids kanamycin resistance. To generate specific amino acid mutations, we used the classical QuikChange site-directed mutagenesis protocol based on polymerase chain reaction (PCR) amplification using *Pfu*Turbo DNA polymerase (Agilent) and two complementary oligonucleotide primers (Microsynth) containing the desired mutation. Following temperature cycling, reactions were treated with *Dpn*I (New England Biolabs) for digestion of parental DNA template. After transformation (XL2-Blue), positive clones were identified and validated by Sanger sequencing. Oligos used for mutagenesis are as follows, with lowercase letters indicating mutated nucleotides:

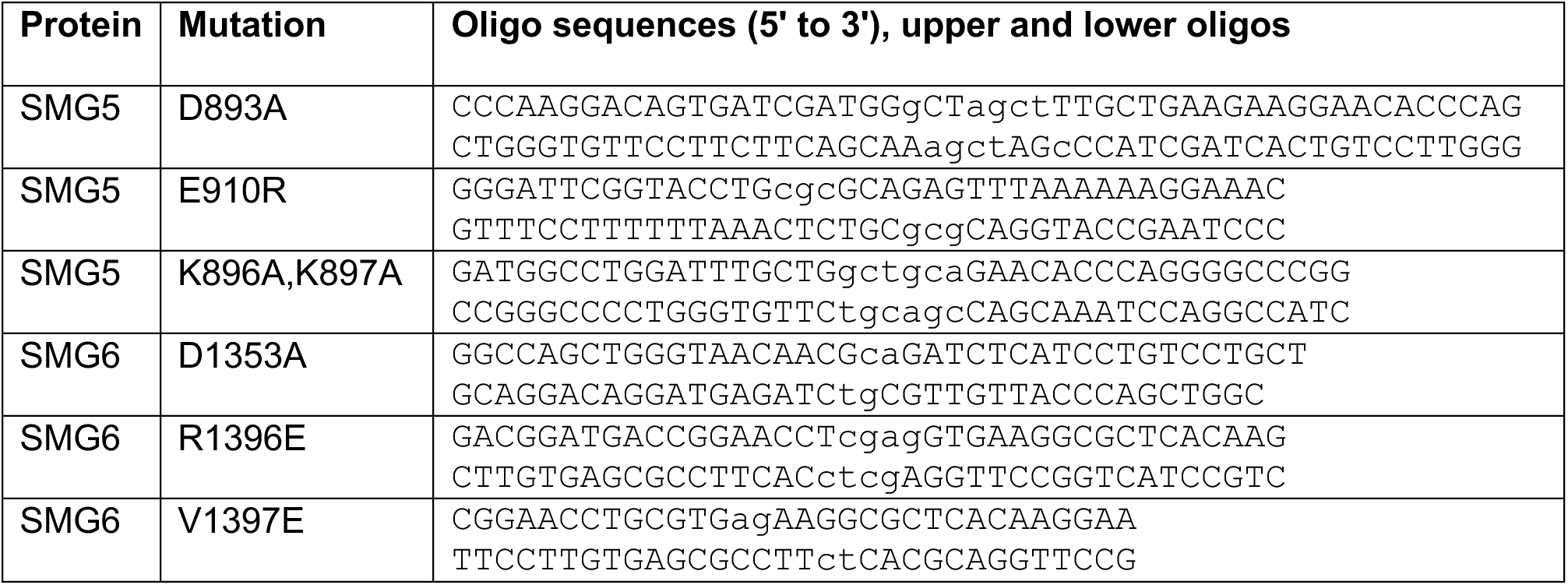

The overall list of DNA constructs used in this study is as follows:

pLIBT7_A_HisHsSMG5(827-C)

pLIBT7_A_HisHsSMG5(827-C)_D893A

pLIBT7_A_HisHsSMG5(827-C)_E910R

pLIBT7_A_HisHsSMG5(827-C)_K896A, K897A

pLIBT7_A_SpyTag-SMG5(827-C)-His

pLIBT7_A_SpyTag-SMG5(827-C)-His_D893A

pLIBT7_A_SpyTag-SMG5(827-C)-His_K896A, K897A

pLIBT7_K_HisGST-TEV-SMG6(1162-C)

pLIBT7_K_HisGST-TEV-SMG6(567-C)

pLIBT7_K_HisGST-TEV-SMG6(567-C)_D1353A

pLIBT7_K_HisGST-TEV-SMG6(567-C)_R1396E

pLIBT7_K_HisGST-TEV-SMG6(567-C)_V1397E

pLIBT7_K_SpyCatcher-SMG6(1162-C)-His

pLIBT7_K_SpyCatcher-SMG6(1162-C)-His_D1353A

pLIBT7_K_SpyCatcher-SMG6(1162-C)-His_V1397E

### Expression, and purification of SMG5 and SMG6 proteins

Pairs of SMG5- and SMG6-expression plasmids were co-transformed into electrocompetent *E. coli* Rosetta(DE3) cells and positive clones were selected on LB plates supplemented with 100 µg/mL ampicillin, 50 µg/mL kanamycin, and 30 µg/mL chloramphenicol. These cells were used to inoculate 1 L cultures in TB medium, and after growth at 37 °C to an OD600 of 0.5-1 the temperature was reduced to 22 °C. Expression was then induced by addition of IPTG to a final concentration of 0.4 mM, and the cultures were left shaking at 22 °C overnight (16 h). Bacteria were collected by centrifugation and the pellet was resuspended in 70 mL of Lysis buffer (50 mM Tris-HCl pH 7.5, 300 mM NaCl, 5% glycerol, 25 mM imidazole) supplemented with 5 mM β-mercaptoethanol and protease inhibitors (Roche). Cells were lysed by sonication on ice (Bandelin sonicator equipped with VS70T-tip) for 12 min total sonication time (40% output, pulsing 1 s ON / 1 s OFF) and the lysate was clarified by centrifugation at 40000× *g* at 4 °C for 30 min. This lysate was then loaded on a 5 mL HisTrap-HP column (Cytiva) equilibrated in Lysis buffer, followed by a 6 column volume (cV) wash with Lysis buffer. Bound material was then eluted with a 10 cV gradient from Lysis buffer to the same buffer supplemented with 500 mM imidazole. His-GST-TEV-SMG6 and His-SMG5 proteins eluted in separate but overlapping peaks, indicating that their interaction was too weak to fully withstand this purification at 300 mM NaCl concentration. Fractions containing the desired proteins were nevertheless pooled, concentrated to a volume of around 500 µL, and injected onto a Superdex200 Increase size-exclusion column equilibrated in SEC buffer (20 mM Tris-HCl pH 7.5, 250 mM NaCl). His-GST-TEV-SMG6 and His-SMG5 proteins eluted in distinct peaks, which were individually pooled and concentrated. The obtained protein solutions were aliquoted, snap-frozen in liquid nitrogen, and stored at -70 °C.

### Expression and purification of SpyTag-SMG5/SpyCatcher-SMG6 pairs

Combinations of the desired plasmid were co-transformed into *E. coli* Rosetta(DE3) and cultures were grown and lysed as described above. After loading on a His-Trap column the dimer eluted slightly separated from excess SpyTag-SMG5 protein, and fractions containing the desired dimer were concentrated to 500 µL followed by Superdex200 Increase size-exclusion chromatography as described above. The dimer eluted well-separated from the remaining excess SpyTag-SMG5 protein, and was concentrated, aliquoted, snap-frozen and stored at -70 °C.

### *In vitro* gel-based nuclease assay

In a reaction volume of 5 µL, 3 µM of 5’-6FAM-dU-U_25_-dU-oligo (IDT) was incubated with 3 µM of recombinant proteins in reaction buffer containing 100 mM NaCl, 10% glycerol, 5 mM MnCl_2_, 25 mM Tris-HCl pH 7.5, 1 mM DTT. In some assays (e.g., to show the specificity of Mn^2+^-based RNA degradation) 1 mM EGTA and/or 30 mM EDTA were added. The reaction mix was then incubated at temperatures indicated in figure legends (42 °C, 37 °C, 10 °C, or 0 °C), for a reaction time of 45 min (or for 0 min for control sample). After incubation, each reaction was first mixed with 5 µL of 2x RNA loading dye (95% formamide, 18 mM EDTA, 0.025% SDS, and Bromophenol Blue), then heated at 95 °C for 5 min, chilled on ice, then loaded on a 10% polyacrylamide gel containing 7 M Urea (Invitrogen, EC68755BOX). Gel electrophoresis was performed at 120 V for 50 min.

### *In vitro* kinetic nuclease assay

Using substrate concentration of 3 µM for the fluorescence-quenched oligo 5’-6FAM-dU-U_25_-dU-3’-BHQ1 (IDT) and 3 µM of recombinant proteins (or RNase-free water as negative control), reactions were carried out in 10 µL and reaction buffer 100 mM NaCl, 10% glycerol, 5 mM MnCl_2_, 1 mM EGTA, 25 mM Tris-HCl pH 7.5, 1 mM DTT. To catch early reaction timepoints reliably, reactions were typically started by adding the proteins as the last component. The reaction was then incubated at 25 °C for 60 min in a flat-bottom black 384-well plate (Corning). Fluorescence signal was measured every minute in a plate reader (Tecan Safire 2). To calculate observed reaction rate constants (*k_obs_*), within one experiment (i.e., on the same plate) raw data were first normalised to 0-100%, followed by non-linear curve fit (single exponential growth to plateau, using GraphPad Prism 10 software) with formula Y=100 * (1 - exp(-*k_obs_* * X)). As we noted that bleaching of signal occurred over time (i.e., 100% signal decays), we also tried curve fit formulas with a bleaching term, but as the obtained *k_obs_* values were near-identical to the simple fits, we report the outcomes from the simple model in this study.

### Structure predictions

AlphaFold3 [28] was used for structure predictions (5 models each) of complexes formed by full length human (*Hs*) SMG5-6-7, *Drosophila melanogaster* (*Dm*) Smg5-6 (no Smg7 ortholog present in flies), or *Caenorhabditis elegans* (*Ce*) SMG-5-6-7 each including U_6_ RNA and 2 Mg^2+^. The highest ranked model of each prediction was used for visualizations with ChimeraX1.10 [39]. Predicted Aligned Error (PAE) plots were generated with PAE viewer [40], while surface conservation scores were calculated using ConSurf [41] with structure models as input and default settings.

### Sequence alignments

Sequences of human SMG5, SMG6 and orthologous proteins were identified via TreeFam [42] or by BLAST search [43] and aligned using the MUSCLE webserver [44] from within JALVIEW [45]. Positional conservation and similarity scores were calculated using the ESPRIPT webserver [46] with default settings.

## CRediT authorship contribution statement

**Enes S. Arpa**: Writing – review & editing, Writing – original draft, Visualization, Validation, Conceptualization, Formal analysis, Investigation, Methodology. **Michael Taschner:** Writing – review & editing, Writing – original draft, Conceptualization, Investigation, Methodology. **Mara De Matos:** Investigation, Methodology. **Stefanie Jonas:** Writing – review & editing, Writing – original draft, Visualization, Funding acquisition, Conceptualization, Methodology, Supervision. **David Gatfield:** Writing – review & editing, Writing – original draft, Visualization, Funding acquisition, Conceptualization, Formal analysis, Supervision, Project administration.

## Acknowledgements

This work was supported by the University of Lausanne and ETH Zurich, Swiss National Science Foundation (SNSF) NCCR RNA & Disease Phase III funding (grant number 205601 to D.G. and S.J.), SNSF Project funding (grant number 212423 to D.G., and 207458 to S.J.), an ERC Consolidator grant funded via SERI (MB22.00064 to S.J.), an EMBO Young Investigators Award (4918 to S.J.).

## References

[1] Peltz SW, Brown AH, Jacobson A. mRNA destabilization triggered by premature translational termination depends on at least three cis-acting sequence elements and one trans-acting factor. Genes & development. 1993;7:1737–54.

[2] Leeds P, Peltz SW, Jacobson A, Culbertson MR. The product of the yeast UPF1 gene is required for rapid turnover of mRNAs containing a premature translational termination codon. Genes & development. 1991;5:2303–14.

[3] Hodgkin J, Papp A, Pulak R, Ambros V, Anderson P. A new kind of informational suppression in the nematode Caenorhabditis elegans. Genetics. 1989;123:301–13.

[4] He F, Jacobson A. Upf1p, Nmd2p, and Upf3p regulate the decapping and exonucleolytic degradation of both nonsense-containing mRNAs and wild-type mRNAs. Molecular and cellular biology. 2001;21:1515–30.

[5] Lejeune F, Li X, Maquat LE. Nonsense-mediated mRNA decay in mammalian cells involves decapping, deadenylating, and exonucleolytic activities. Molecular cell. 2003;12:675–87.

[6] Karousis ED, Muhlemann O. The broader sense of nonsense. Trends in biochemical sciences. 2022;47:921–35.

[7] Muhlrad D, Parker R. Premature translational termination triggers mRNA decapping. Nature. 1994;370:578–81.

[8] Mitchell P, Tollervey D. An NMD pathway in yeast involving accelerated deadenylation and exosome-mediated 3’-->5’ degradation. Molecular cell. 2003;11:1405–13.

[9] Gatfield D, Izaurralde E. Nonsense-mediated messenger RNA decay is initiated by endonucleolytic cleavage in Drosophila. Nature. 2004;429:575–8.

[10] Glavan F, Behm-Ansmant I, Izaurralde E, Conti E. Structures of the PIN domains of SMG6 and SMG5 reveal a nuclease within the mRNA surveillance complex. The EMBO journal. 2006;25:5117–25.

[11] Huntzinger E, Kashima I, Fauser M, Sauliere J, Izaurralde E. SMG6 is the catalytic endonuclease that cleaves mRNAs containing nonsense codons in metazoan. Rna. 2008;14:2609–17.

[12] Eberle AB, Lykke-Andersen S, Muhlemann O, Jensen TH. SMG6 promotes endonucleolytic cleavage of nonsense mRNA in human cells. Nature structural & molecular biology. 2009;16:49–55.

[13] Chakrabarti S, Bonneau F, Schussler S, Eppinger E, Conti E. Phospho-dependent and phospho-independent interactions of the helicase UPF1 with the NMD factors SMG5-SMG7 and SMG6. Nucleic acids research. 2014;42:9447–60.

[14] Barbarin-Bocahu I, Ulryck N, Rigobert A, Ruiz Gutierrez N, Decourty L, Raji M, et al. Structure of the Nmd4-Upf1 complex supports conservation of the nonsense-mediated mRNA decay pathway between yeast and humans. PLoS biology. 2024;22:e3002821.

[15] Langer LM, Kurscheidt K, Basquin J, Bonneau F, Iermak I, Basquin C, et al. UPF1 helicase orchestrates mutually exclusive interactions with the SMG6 endonuclease and UPF2. Nucleic acids research. 2024;52:6036–48.

[16] Okada -Katsuhata Y, Yamashita A, Kutsuzawa K, Izumi N, Hirahara F, Ohno S. N- and C-terminal Upf1 phosphorylations create binding platforms for SMG-6 and SMG-5:SMG-7 during NMD. Nucleic acids research. 2012;40:1251–66.

[17] Unterholzner L, Izaurralde E. SMG7 acts as a molecular link between mRNA surveillance and mRNA decay. Molecular cell. 2004;16:587–96.

[18] Jonas S, Weichenrieder O, Izaurralde E. An unusual arrangement of two 14-3-3-like domains in the SMG5-SMG7 heterodimer is required for efficient nonsense-mediated mRNA decay. Genes & development. 2013;27:211–25.

[19] Loh B, Jonas S, Izaurralde E. The SMG5-SMG7 heterodimer directly recruits the CCR4-NOT deadenylase complex to mRNAs containing nonsense codons via interaction with POP2. Genes & development. 2013;27:2125–38.

[20] Colombo M, Karousis ED, Bourquin J, Bruggmann R, Muhlemann O. Transcriptome-wide identification of NMD-targeted human mRNAs reveals extensive redundancy between SMG6-and SMG7-mediated degradation pathways. Rna. 2017;23:189–201.

[21] Boehm V, Kueckelmann S, Gerbracht JV, Kallabis S, Britto-Borges T, Altmuller J, et al. SMG5-SMG7 authorize nonsense-mediated mRNA decay by enabling SMG6 endonucleolytic activity. Nature communications. 2021;12:3965.

[22] Huth M, Santini L, Galimberti E, Ramesmayer J, Titz-Teixeira F, Sehlke R, et al. NMD is required for timely cell fate transitions by fine-tuning gene expression and regulating translation. Genes & development. 2022;36:348–67.

[23] Viscardi MJ, Sehgal E, Arribere JA. Endonucleolytic cleavage is the primary mechanism of decay elicited by C. elegans nonsense-mediated mRNA decay. Genome research. 2025;35:1337–48.

[24] Senissar M, Manav MC, Brodersen DE. Structural conservation of the PIN domain active site across all domains of life. Protein Sci. 2017;26:1474–92.

[25] Matelska D, Steczkiewicz K, Ginalski K. Comprehensive classification of the PIN domain-like superfamily. Nucleic acids research. 2017;45:6995–7020.

[26] Arcus VL, McKenzie JL, Robson J, Cook GM. The PIN-domain ribonucleases and the prokaryotic VapBC toxin-antitoxin array. Protein Eng Des Sel. 2011;24:33–40.

[27] Katsioudi G, Dreos R, Arpa ES, Gaspari S, Liechti A, Sato M, et al. A conditional Smg6 mutant mouse model reveals circadian clock regulation through the nonsense-mediated mRNA decay pathway. Sci Adv. 2023;9:eade2828.

[28] Abramson J, Adler J, Dunger J, Evans R, Green T, Pritzel A, et al. Accurate structure prediction of biomolecular interactions with AlphaFold 3. Nature. 2024;630:493–500.

[29] Gatfield D, Unterholzner L, Ciccarelli FD, Bork P, Izaurralde E. Nonsense-mediated mRNA decay in Drosophila: at the intersection of the yeast and mammalian pathways. The EMBO journal. 2003;22:3960–70.

[30] Das U, Pogenberg V, Subhramanyam UK, Wilmanns M, Gourinath S, Srinivasan A. Crystal structure of the VapBC-15 complex from Mycobacterium tuberculosis reveals a two-metal ion dependent PIN-domain ribonuclease and a variable mode of toxin-antitoxin assembly. J Struct Biol. 2014;188:249–58.

[31] Zakeri B, Fierer JO, Celik E, Chittock EC, Schwarz-Linek U, Moy VT, et al. Peptide tag forming a rapid covalent bond to a protein, through engineering a bacterial adhesin. Proceedings of the National Academy of Sciences of the United States of America. 2012;109:E690–7.

[32] Nicholson P, Josi C, Kurosawa H, Yamashita A, Muhlemann O. A novel phosphorylation-independent interaction between SMG6 and UPF1 is essential for human NMD. Nucleic acids research. 2014;42:9217–35.

[33] Fukuhara N, Ebert J, Unterholzner L, Lindner D, Izaurralde E, Conti E. SMG7 is a 14-3-3-like adaptor in the nonsense-mediated mRNA decay pathway. Molecular cell. 2005;17:537–47.

[34] Durand S, Franks TM, Lykke-Andersen J. Hyperphosphorylation amplifies UPF1 activity to resolve stalls in nonsense-mediated mRNA decay. Nature communications. 2016;7:12434.

[35] Boehm V, Haberman N, Ottens F, Ule J, Gehring NH. 3’ UTR length and messenger ribonucleoprotein composition determine endocleavage efficiencies at termination codons. Cell reports. 2014;9:555–68.

[36] Cook GM, Robson JR, Frampton RA, McKenzie J, Przybilski R, Fineran PC, et al. Ribonucleases in bacterial toxin-antitoxin systems. Biochimica et biophysica acta. 2013;1829:523–31.

[37] Wang F, Zhao P, Bi X, Zheng R, Tian X, Xu J, et al. Cyclic tetraadenylate binding induces dimerization of protein dimers to activate a CRISPR-associated PIN nuclease. Nucleic acids research. 2025;53.

[38] Kashima I, Jonas S, Jayachandran U, Buchwald G, Conti E, Lupas AN, et al. SMG6 interacts with the exon junction complex via two conserved EJC-binding motifs (EBMs) required for nonsense-mediated mRNA decay. Genes & development. 2010;24:2440–50.

[39] Meng EC, Goddard TD, Pettersen EF, Couch GS, Pearson ZJ, Morris JH, et al. UCSF ChimeraX: Tools for structure building and analysis. Protein Sci. 2023;32:e4792.

[40] Elfmann C, Stulke J. PAE viewer: a webserver for the interactive visualization of the predicted aligned error for multimer structure predictions and crosslinks. Nucleic acids research. 2023;51:W404–W10.

[41] Yariv B, Yariv E, Kessel A, Masrati G, Chorin AB, Martz E, et al. Using evolutionary data to make sense of macromolecules with a “face-lifted” ConSurf. Protein Sci. 2023;32:e4582.

[42] Ruan J, Li H, Chen Z, Coghlan A, Coin LJ, Guo Y, et al. TreeFam: 2008 Update. Nucleic acids research. 2008;36:D735–40.

[43] Altschul SF, Madden TL, Schaffer AA, Zhang J, Zhang Z, Miller W, et al. Gapped BLAST and PSI-BLAST: a new generation of protein database search programs. Nucleic acids research. 1997;25:3389–402.

[44] Edgar RC. MUSCLE: multiple sequence alignment with high accuracy and high throughput. Nucleic acids research. 2004;32:1792–7.

[45] Waterhouse AM, Procter JB, Martin DM, Clamp M, Barton GJ. Jalview Version 2--a multiple sequence alignment editor and analysis workbench. Bioinformatics. 2009;25:1189–91.

[46] Robert X, Gouet P. Deciphering key features in protein structures with the new ENDscript server. Nucleic acids research. 2014;42:W320–4.

