## Supplemental Figures S1-S5 for "Active site assembly by SMG5 as a mechanism for SMG6 endonuclease licensing in nonsense-mediated mRNA decay"

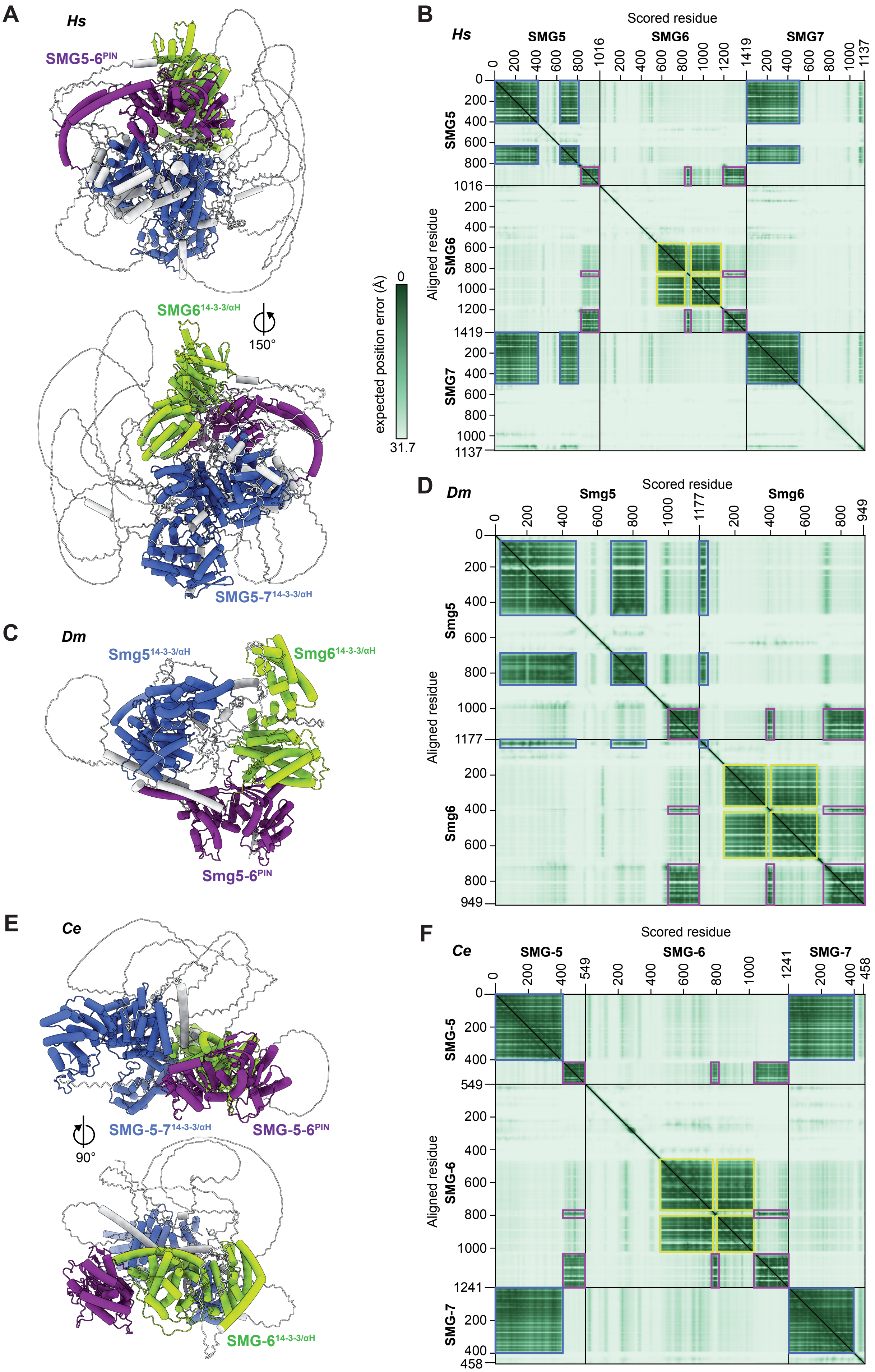
A
Hs
SMG5-6PIN
SMG614-3-3/αH
150°
SMG5-714-3-3/αH
B
Hs
Scored residue
SMG5
SMG6
SMG7
Aligned residue
expected position error (Å)
0
31.7
C
Dm
Smg514-3-3/αH
Smg614-3-3/αH
Smg5-6PIN
D
Dm
Scored residue
Smg5
Smg6
Aligned residue
E
Ce
SMG-5-714-3-3/αH
SMG-5-6PIN
90°
SMG-614-3-3/αH
F
Ce
Scored residue
SMG-5
SMG-6
SMG-7
Aligned residue

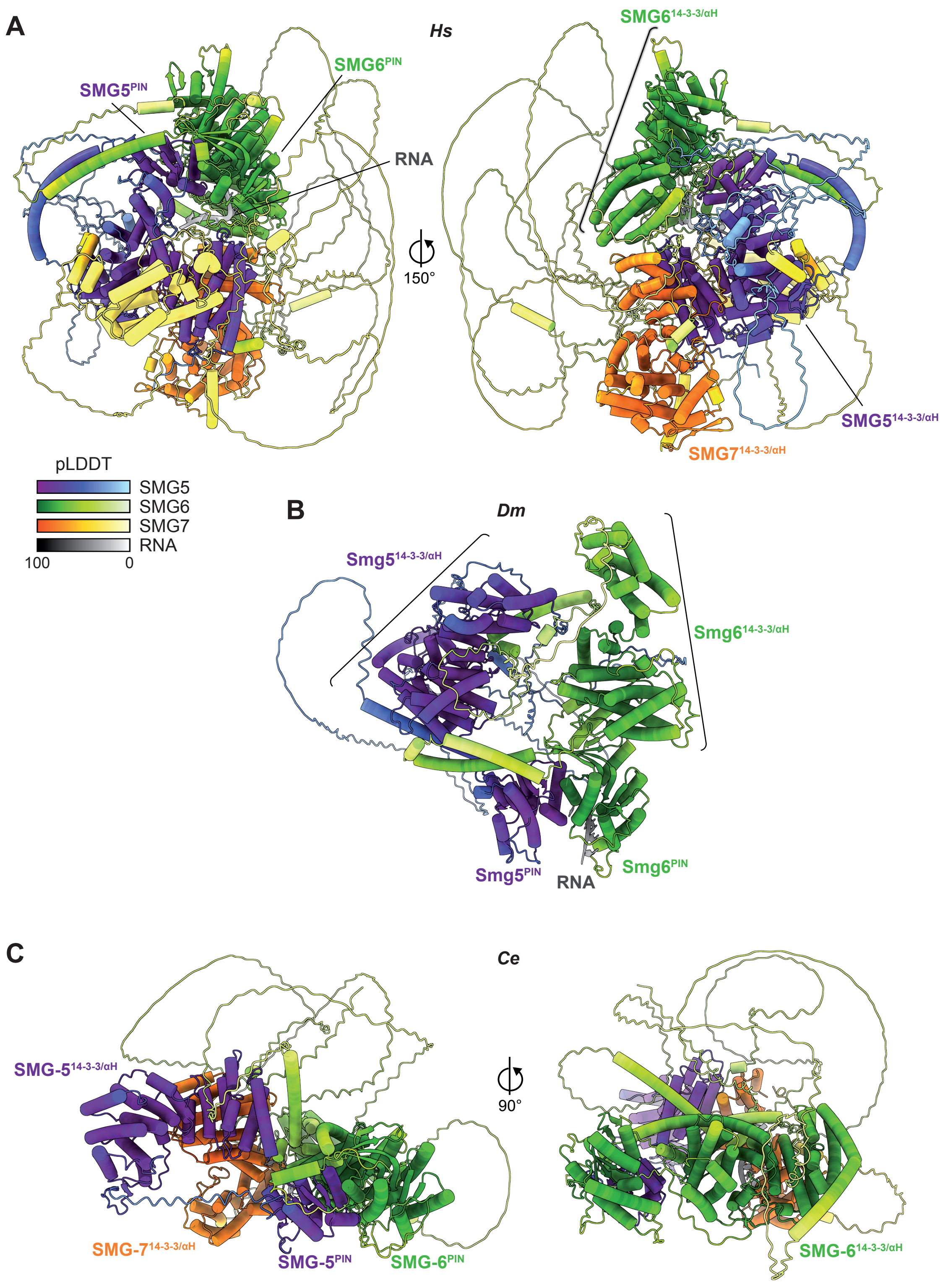

A
Hs
SMG5PIN
SMG6PIN
RNA
150°
SMG614-3-3/αH
SMG514-3-3/αH
SMG714-3-3/αH
pLDDT
SMG5
SMG6
SMG7
RNA
100
0
B
Dm
Smg514-3-3/αH
Smg614-3-3/αH
Smg5PIN
RNA
Smg6PIN
C
Ce
SMG-514-3-3/αH
90°
SMG-714-3-3/αH
SMG-5PIN
SMG-6PIN
SMG-614-3-3/αH

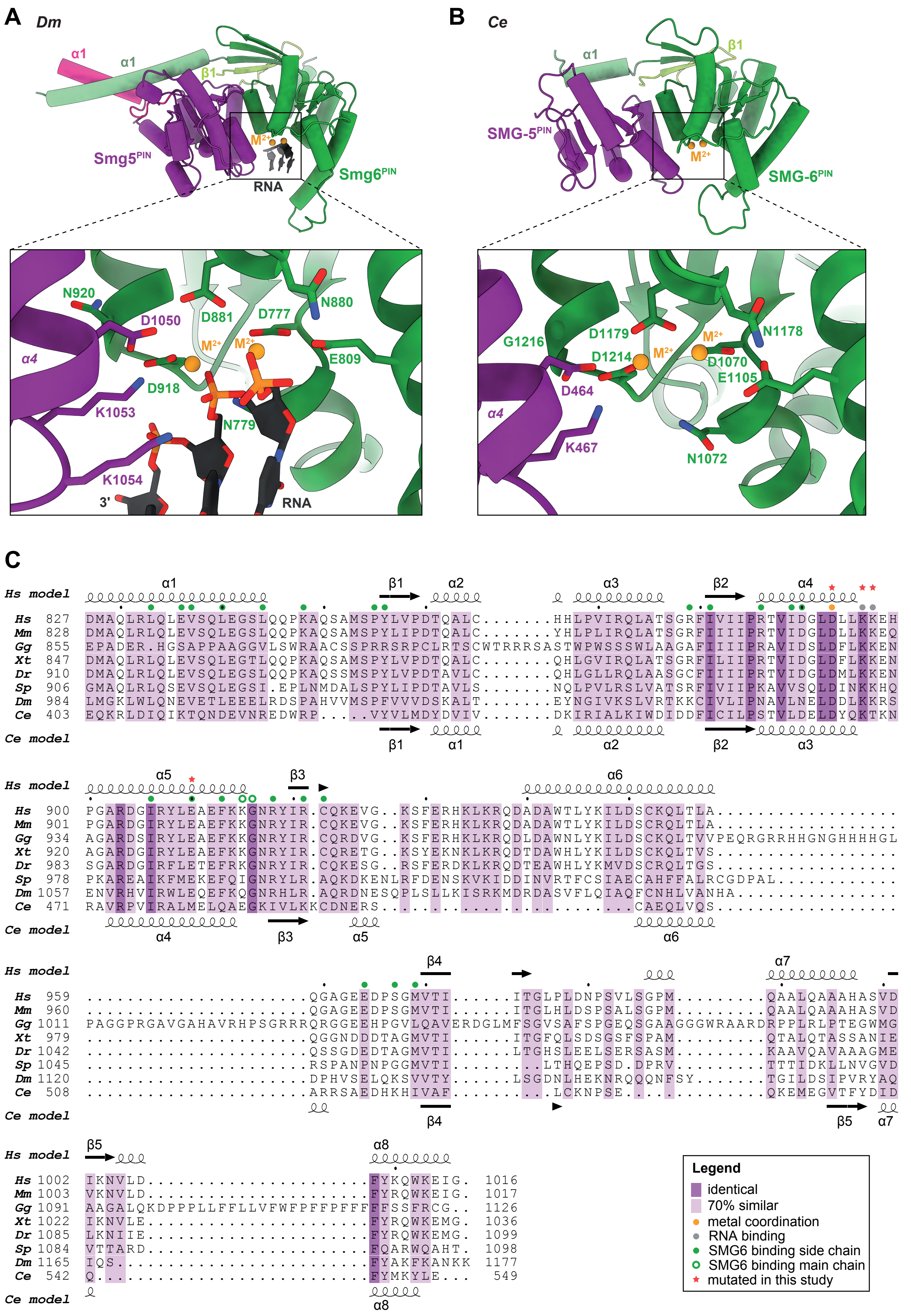
A Dm
α1
α1
β1
Smg5PIN
M2+
RNA
Smg6PIN
N920
D1050
D881
D777
N880
M2+
M2+
α4
E809
D918
K1053
N779
K1054
3'
RNA
B Ce
α1
β1
SMG-5PIN
M2+
SMG-6PIN
D1179
M2+
N1178
G1216
D1214
M2+
D1070
E1105
D464
α4
K467
N1072
C
Hs model
Hs 827 DMAQLRLQLEVSQLEGSLQQPKAQSAMSPYLVPDTQALC.......HHLPVIRQLATSGRFIVIIPRTVIDGLDLLKKEH
Mm 828 DMAQLRLQLEVSQLEGSLQQPKAQSAMSPYLIPDTQALC.......YHLPLIRQLATSGRFIIIIPRTVIDGLDLLKKEQ
Gg 855 EPADER.HGSAPPAAGGVLSWRAACSSPRRSRPCLRTSCWTRRRSASTWPWSSSWLAAGAFIIIIPRAVIDSLDFLKKEN
Xt 847 DMAQLRLQLEVSQLEGTLQQPKAQSAMSPYLVPDTQALC.......QHLGVIRQLATSGRFILIIPRTVIDGLDFLKKEN
Dr 910 DMAQLRLQLEVSQLEGSLQQPKAQSSMSPYLVPDTAVLC.......QHLGLLRQLAASGCFIIIIPRTVIDGLDMLKKEN
Sp 906 GMAQLRLQSEVSQLEGSI.EPLNMDALSPYLIPDTAVLS.......NQLPVLRSLVATSRFIVIIPKAVVSQLDINKKHQ
Dm 984 LMGKLWLQNEVETLEEELRDSPAHVVMSPFVVVDSKALT.......EYNGIVKSLVRTKKCIVLIPNAVLNELDDLKKRS
Ce 403 EQKRLDIQIKTQNDEVNREDWRP....VYVLMDYDVIV.......DKIRIALKIWDIDDFICILPSTVLDELDYQKTKN
Ce model
α1 β1 α2 α3 β2 α4
β1 α1 α2 β2 α3
Hs model
α5 β3 α6
Hs 900 PGARDGIRYLEAEFKKGNRYIR.CQKEVG..KSFERHKLKRQDADAWTLYKILDSCKQLTLA.................
Mm 901 PGARDGIRYLEAEFKKGNRYIR.CQKEVG..KSFERHKLKRQDADAWTLYKILDSCRQLTLA.................
Gg 934 AGARDSIRYLEAEFKKGNRYIR.CQRDVG..KSFERHKLKRQDLDAWNLYKILDSCKQLTVVPEQRGRRHHGNGHHHHGL
Xt 920 AGARDGIRYLEAEFKKGNRYIR.CQRETG..KSYEKNKLKRQDTDAWHLYKILDSCKQLTVS.................
Dr 983 SGARDGIRFLETEFRKGNRYIR.CQKESG..RSFERDKLKRQDTEAWHLYKMVDSCRQLTGS.................
Sp 978 PKAREAIKFMEKEFQIGNRYLR.AQKDKENLRFDENSKVKIDDINVRTFCSIAECAHFFALRCGDPAL...........
Dm 1057 ENVRHVIRWLEQEFKQGNRHLR.AQRDNESQPLSLLKISRKMDRDASVFLQIAQFCNHLVANHA...............
Ce 471 RAVRPVIRALMELQAEGKIVLKKCDNERS........................CAEQLVQS.................
Ce model
α4 β3 α5 α6
Hs model
β4 α7
Hs 959 ...........................QGAGEEDPSGMVTI......ITGLPLDNPSVLSGPM.........QAALQAAAHASVD
Mm 960 ...........................QGAGEEDPSGMVTI......ITGLHLDSPSALSGPM.........QAALQAAAHASVD
Gg 1011 PAGGPRGAVGAHAVRHPSGRRRQRGGEEHPGVLQAVERDGLMFSGVSAFSPGEQSGAAGGGWRAARDRPPLRLPTEGWMG
Xt 979 ...........................QGGNDDDTAGMVTI......ITGFQLSDSGSFSPAM.........QTALQTASSANIE
Dr 1042 ...........................QSSGDEDTAGMVTI......LTGHSLEELSERSASM.........KAAVQAVAAAGME
Sp 1045 ...........................RSPANPNPGGMVTI........LTHQEPSD.DPRV.........TTTIDKLLNVGVD
Dm 1120 ...........................DPHVSELQKSVVTY......LSGDNLHEKNRQQQNFSY.......TGILDSIPVRYAQ
Ce 508 ...........................ARRSAEDHKHIVAF.........LCKNPSE.................QKEMEGVTFYDID
Ce model
β4 β5 α7
Hs model
β5 α8
Hs 1002 IKNVLD..........................FYKQWKEIG. 1016
Mm 1003 VKNVLD..........................FYRQWKEIG. 1017
Gg 1091 AAGALQKDPPPLLFFLLVFWFPFFPFFFFFFSSFRCG.. 1126
Xt 1022 IKNVLE..........................FYRQWKEMG. 1036
Dr 1085 LKNIIE..........................FYRQWKEMG. 1099
Sp 1084 VTTARD..........................FQARWQAHT. 1098
Dm 1165 IQS.............................FYAKFKANKK 1177
Ce 542 Q...............................FYMKYLE... 549
Ce model
α8
Legend
identical
70% similar
metal coordination
RNA binding
SMG6 binding side chain
SMG6 binding main chain
mutated in this study

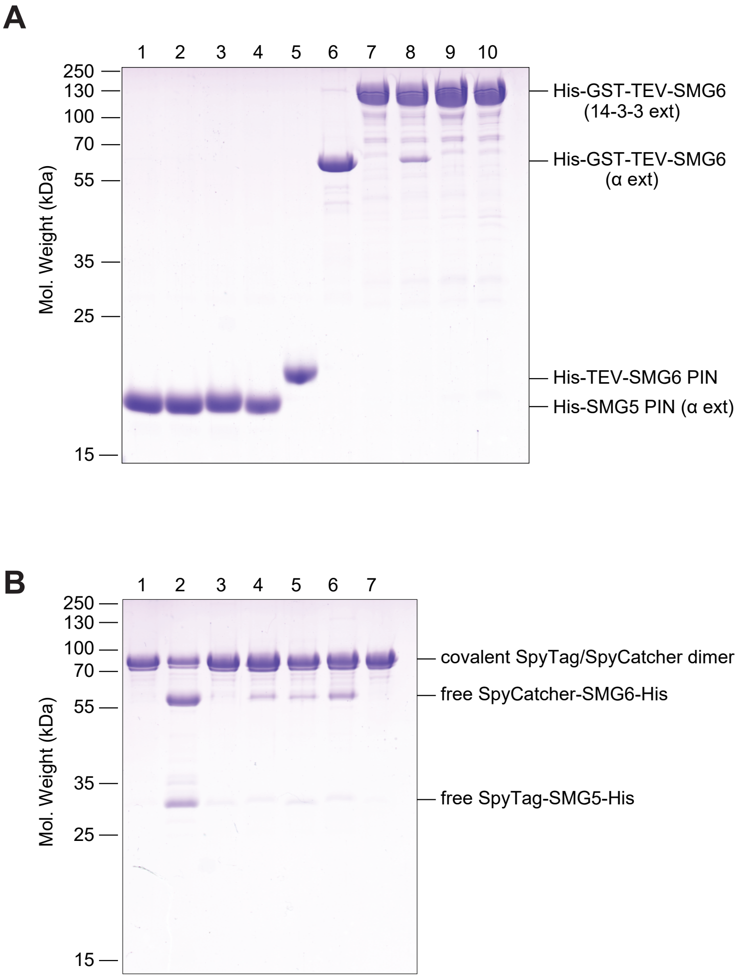


1: His-SMG5 PIN(α ext); wt
2: His-SMG5 PIN(α ext); D893A
3: His-SMG5 PIN(α ext); K896A/K897A
4: His-SMG5 PIN(α ext); E910R
5: His-TEV-SMG6 PIN; wt
6: His-GST-TEV-SMG6(α ext); wt
7: His-GST-TEV-SMG6(14-3-3 ext); wt
8: His-GST-TEV-SMG6(14-3-3 ext); D1353A
9: His-GST-TEV-SMG6(14-3-3 ext); R1396E
10: His-GST-TEV-SMG6(14-3-3 ext); V1397E

1: SpyTag-SMG5PIN(α ext)-His/
SpyCatcher-SMG6(α ext)-His
2: SpyTag-SMG5PIN(α ext)-His /
SpyCatcher-SMG6(α ext; V1397E)-His
3: SpyTag-SMG5PIN(α ext)-His /
SpyCatcher-SMG6(α ext; D1353A)-His
4: SpyTag-SMG5PIN(α ext; D893A)-His /
SpyCatcher-SMG6(α ext)-His
5: SpyTag-SMG5PIN(α ext; D893A)-His /
SpyCatcher-SMG6(α ext;D1353A)-His
6: SpyTag-SMG5PIN(α ext; K896A;K897A)-His /
SpyCatcher-SMG6(α ext)-His
7: SpyTag-SMG5PIN(α ext; K896A;K897A)-His /
SpyCatcher-SMG6(α ext; D1353A)-His

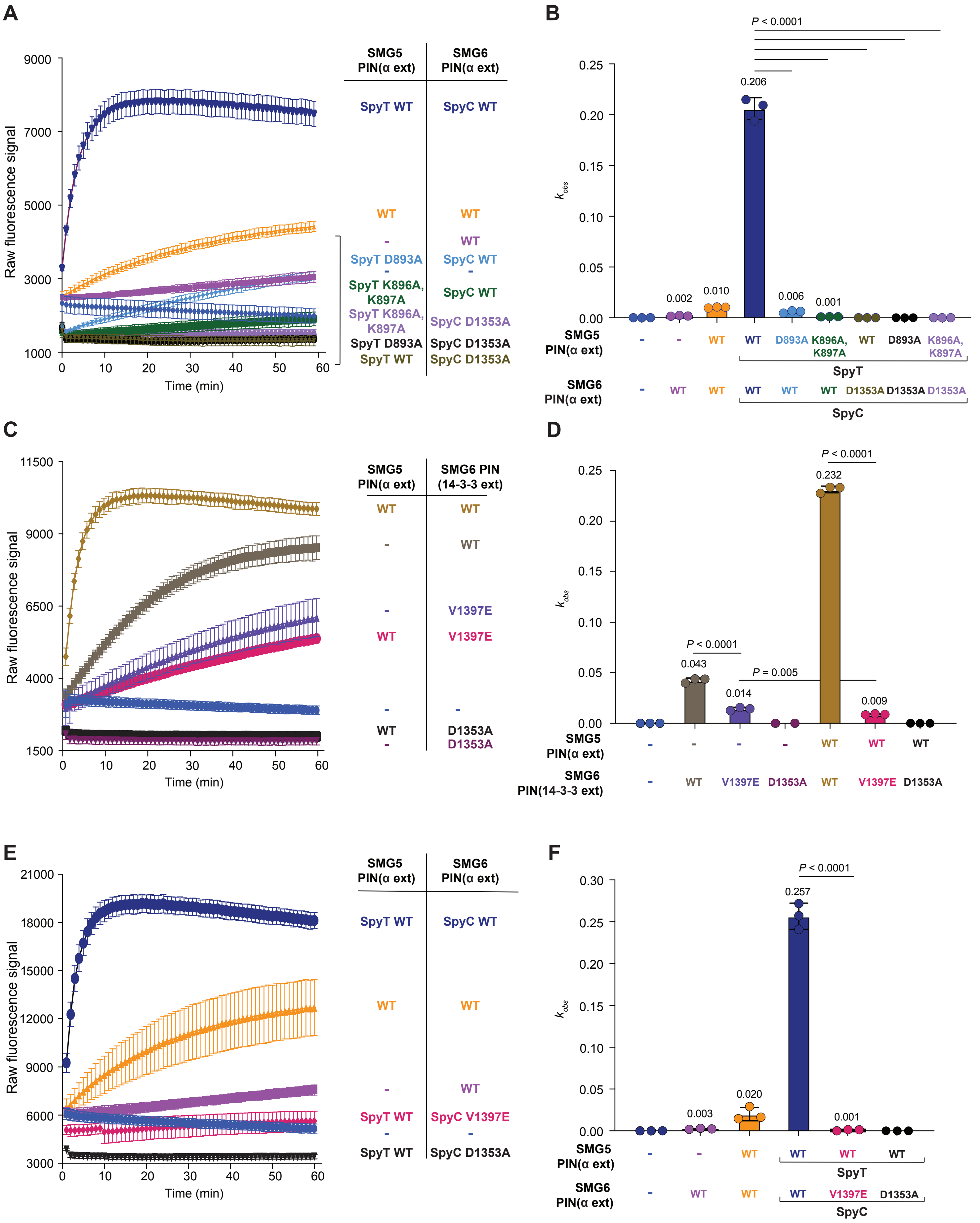
A
B
C
D
E
F
Raw fluorescence signal
Time (min)
$k_{obs}$
SMG5 PIN(α ext)
SMG6 PIN(α ext)
SMG6 PIN (14-3-3 ext)
PIN(14-3-3 ext)
SpyT WT
SpyC WT
WT
-
SpyT D893A
SpyT K896A, K897A
SpyC D1353A
SpyC V1397E
V1397E
D1353A
D893A
K896A, K897A
SpyT
SpyC
P < 0.0001
P = 0.005
0.206
0.002
0.010
0.006
0.001
0.232
0.043
0.014
0.009
0.257
0.003
0.020
